# Cytotoxicity-based High-throughput Screening System for CAR T Cell

**DOI:** 10.64898/2026.03.31.715247

**Authors:** Atsushi Okuma, Yoshihito Ishida, Takuya Miura-Yamashita, Taketo Kawara, Daisuke Ito, Kei Yoshida, Satoru Mimura, Yuki Nakao, Tatsuki Iwamoto, Shoji Hisada, Shizu Takeda

## Abstract

Some chimeric antigen receptor (CAR) T cell therapies have shown strong clinical efficacy, yet systematic screening of new CAR designs remains constrained by labor-intensive, low-throughput evaluation methods. To address this limitation, we developed a cytotoxicity-centered, high-throughput screening platform that integrates single-cell pooled screening with fully automated arrayed screening enabling both large-scale library handling and quantitative functional resolution for systematic CAR design exploration. Using a mutation-based CAR design approach guided by protein fitness prediction, we generated a 4-1BB–based CAR library with approximately 10⁸ theoretical variants while minimizing the prevalence of low-activity designs. In pooled screening, CAR T cells were evaluated at the single-cell level based on cytotoxicity and proliferation, enabling rapid enrichment of high-performing variants from a highly diverse library. Subsequent automated arrayed screening quantitatively measured cytotoxicity with high reproducibility, providing high-resolution functional data suitable for comparative ranking. Selected CAR variants demonstrated superior antitumor efficacy in a leukemia xenograft model compared with a template CAR. Furthermore, systematic analysis of mutation sites from an enhanced CAR variant identified essential mutation combinations underlying functional enhancement. Together, this study establishes a cytotoxicity-focused screening framework that provides a robust approach for optimizing CAR architectures and accelerating the development of CAR T-cell therapies.

## INTRODUCTION

Chimeric antigen receptor (CAR) T cell therapies have demonstrated significant clinical efficacy in the treatment of B cell malignancies and multiple myeloma. The development of CAR T cell therapeutics was spearheaded by the CD19-targeted CAR T cell products axicabtagene ciloleucel (axi-cel) and tisagenlecleucel (tisa-cel), both of which were approved in 2017. Consequently, these two products have been frequently compared, as each CAR design possessing unique characteristics (Bachy et al., 2022; Kawalekar et al., 2016; Long et al., 2015; Yu et al., 2025). Currently, challenges remain in CAR T cell therapy including recurrence, adverse events, on-target off-tumor effect, and the microenvironment of solid tumors. To address these challenges, there is a growing need to explore CAR designs that extend beyond a limited set of domains currently in clinical use, which primarily includes components derived from IgG4, CD8α, 4-1BB, CD28 and CD3ζ (Labanieh and Mackall, 2023; Singh and Maus, 2023). This search for solutions has motivated the engineering of novel CAR designs that provide unique activation properties.

The development of new CAR architecture remains challenges because conventional CAR T cell generation and evaluation processes are time-consuming and costly. While pooled screening strategies have greatly reduced experimental labor, most rely on surrogate readouts such as activation markers, cytokine production, and/or *in vitro* fitness (Castellanos-Rueda et al., 2025, 2022; Daniels et al., 2022; Goodman et al., 2022; Gordon et al., 2022; Rios et al., 2023), whose ability to predict *in vivo* efficacy remains uncertain. To move beyond such surrogate readouts, single-cell transcriptomic profiling has been increasingly adopted to characterize CAR T cell states in pooled libraries (Castellanos-Rueda et al., 2025, 2022; Perez et al., 2025). However, the scalability of single-cell RNA-seq–based approaches is inherently limited, restricting their applicability to libraries containing only dozens of CAR variants.

In contrast to pooled approaches, arrayed screening strategies enable direct and quantitative evaluation of CAR T cell function using bulk populations, thereby providing high reproducibility and assay resolution. Indeed, arrayed assays have been successfully used to generate high-quality datasets suitable for comparative ranking and machine-learning applications (Daniels et al., 2022). However, the throughput of arrayed screening is inherently limited by the need to individually construct, produce, and evaluate each CAR design, making it impractical for exploring highly diverse CAR libraries. As a result, arrayed screening alone cannot efficiently address the vast design space required for systematic CAR optimization.

Regarding CAR design strategies, CAR libraries that incorporate selected intracellular domains have been employed for pooled screening, primarily due to the impact of the differences between CD28 and 4-1BB costimulatory domains on clinical outcome and phenotypic characteristics of approved CD19 CARs (Bachy et al., 2022; Cappell and Kochenderfer, 2021; Kawalekar et al., 2016; Long et al., 2015; Yu et al., 2025). However, this strategy lacks diversity in CAR libraries, with limited alternatives to CD28 or 4-1BB (Castellanos-Rueda et al., 2022; Goodman et al., 2022; Rios et al., 2023). To increase diversity, CAR libraries have also adopted combinations of multiple costimulatory domains (Gordon et al., 2022; Perez et al., 2025), signaling motifs (Daniels et al., 2022), or other domains such as hinge domain (Rios et al., 2023). Alternatively, the point mutation strategy can be used to design highly diverse CAR architectures as well. Indeed, one or more point mutations on CD8α hinge (Chen et al., 2022), CD28 hinge (Folimonova et al., 2024), 4-1BB intracellular domain (ICD) (Li et al., 2020), CD28 ICD (Boucher et al., 2021; Guedan et al., 2020), or CD3ζ ICD (Feucht et al., 2019) have been reported to enhance CAR T cell efficacy. Together, these studies highlight the potential of mutation-based strategies to expand CAR design diversity, while underscoring the need for screening platforms capable of handling such diversity.

Here, we developed a CAR T cell screening platform that integrates cytotoxicity-based pooled screening with fully automated arrayed screening to efficiently identify effective CAR T cells from highly diverse CAR libraries. Using this platform, we introduce a mutation-based CAR design strategy, termed multi-domain protein designer (MDPD), which enables the generation of CAR libraries with theoretical diversity approximately 10⁸ variants. Application of this integrated screening workflow successfully enriched CAR variants with enhanced cytotoxicity and superior *in vivo* efficacy compared with a template 4-1BB–based CAR and further enabled the identification of essential mutation combinations underlying functional enhancement.

## RESULTS

### Screening flow overview

To enable cytotoxicity-based evaluation of highly diverse CAR libraries while maintaining quantitative resolution, we developed a stepwise high-throughput screening (HTS) platform that integrates cytotoxicity-based pooled screening with automated arrayed screening (Figure 1A). Initially, we generated pooled CAR T cells and reduced library diversity through flow cytometry-based prescreening followed by a high-throughput single-cell assay. After the pooled screening, we selected highly cytotoxic and higher-growing CAR T cells and analyzed CAR sequences of the selected CAR T cells. We re-synthesized selected CAR-coding sequences and reconstructed CAR vectors. For an arrayed screening, we performed the whole process from viral vector generation to cytotoxicity assay in an automated manner. Selected CAR T cells exhibiting *in vitro* higher cytotoxicity were further evaluated *in vivo* using a leukemia xenograft model.

**Figure 1:**
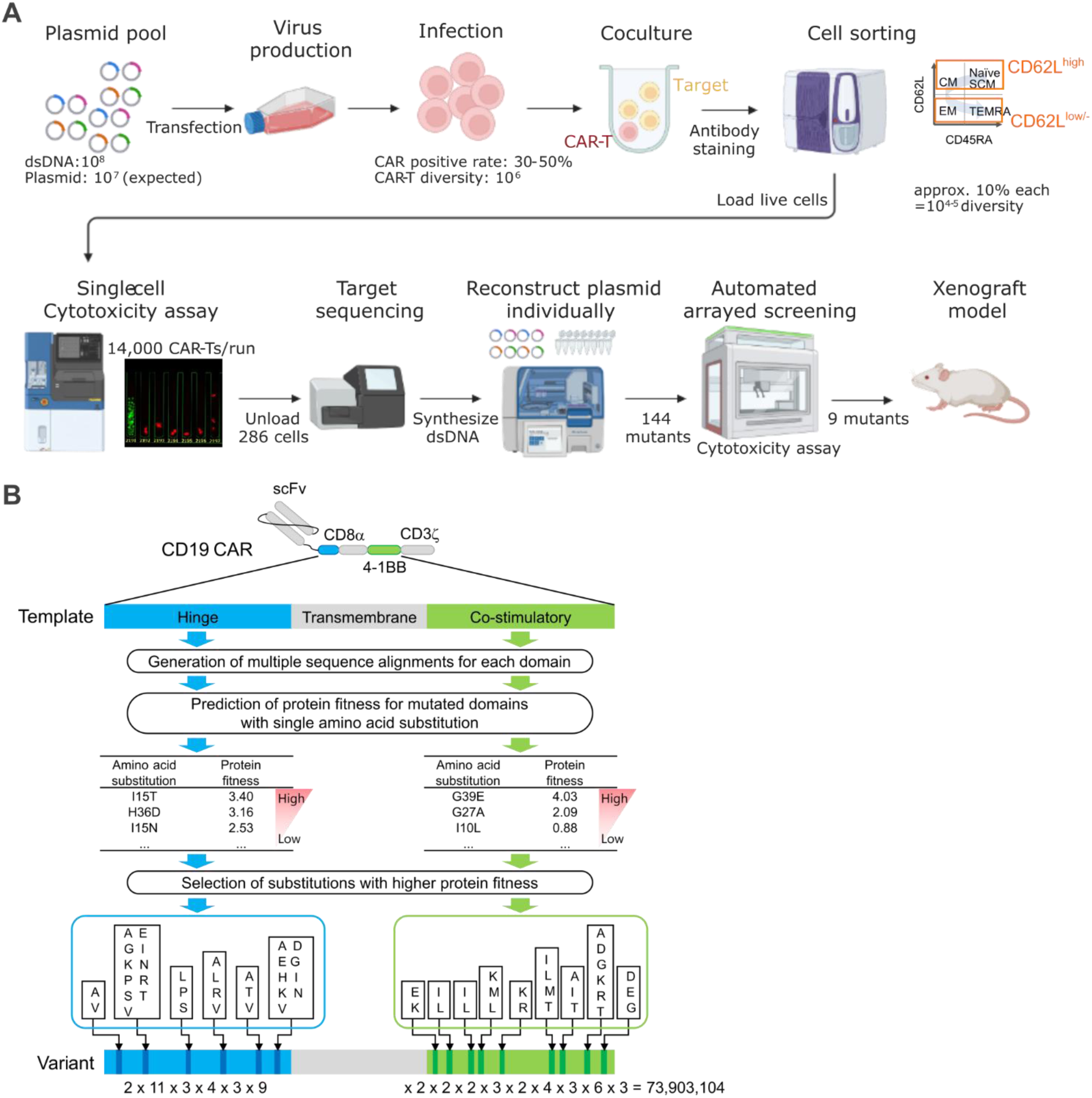
Cytotoxicity-based high-throughput CAR T cell screening platform. (A) Overview of wet screening part of the DesignCell platform. Created with BioRender.com. Human primary T cells were transduced with a lentivirus vector pool coding CAR variants with a diversity of 10^6^-10^7^. The CAR T cell pool was stimulated with CD19-expressing K562 cells (target) and then was sorted out CAR^+^/CD62L^high^ population and CAR^+^/CD62L^low/-^ population. Both populations were loaded into Beacon and cocultured with GFP-expressing Nalm6. CAR T cells with both 100% killing and high growth were unloaded and analyzed CAR sequences with NGS. From the NGS data, 144 plasmids coding CAR variants were reconstructed individually, and cytotoxicity assays were performed using an automated system (see Figure 3A). Based on the quantified cytotoxicity, selected 9 variants were tested *in vivo* mouse model. (B) Schematics of design strategy of a CAR mutation library.

### Generation of CAR sequences

A major challenge in mutation-based CAR library design is to avoid an overwhelming fraction of low-activity or non-functional variants. We applied MDPD to generate a combinatorial mutant library of CARs to identify variants with enhanced cytotoxicity (Figure 1B). MDPD leverages the modular architecture of CARs to expand sequence diversity by introducing amino acid substitutions within individual domains. Using a well-characterized CD19 CAR as a template composed of the FMC63 scFv, a CD8α hinge and transmembrane domain, a 4-1BB costimulatory domain, and CD3ζ (hereafter FMC63/8/8/BBz), we introduced amino acid substitutions into the hinge and costimulatory domains. To minimize the emergence of loss-of-function or very low-activity variants, we prioritized substitutions with higher predicted protein fitness, as estimated by EVmutation (Hopf et al., 2017). We synthesized a double-stranded DNA library encoding all combinations of the selected substitutions, yielding a theoretical diversity of 73,903,104 variants.

### Preparation of CD19 CAR T cell pool

We transduced human primary CD8^+^ T cells with a lentiviral vector pool carrying the CD19 CAR variants designed by MDPD. To reduce the probability of a single cell being infected by multiple vectors, transduction efficiency was adjusted to 30-50% with dilution of the viral vector pool. We estimated the diversity of CAR sequences in the CAR T cell pool by sampling 10^5^ cells and sequencing the variable region of CAR-encoding sequences (Figure S1A). The CD19 CAR T cell pool was activated with CD19-expressing K562 cells for 2 days and stained with antibodies against CD45RA, CD62L, and CD39 to measure T-cell differentiation status. We sorted out CAR^+^ (GFP^+^)/CD39^+^/CD62L^high^ population (naïve, stem cell memory, and central memory) and CAR^+^/CD39^+^/CD62L^low^ population (effector memory and effector) for subsequent screening steps (Figure 1A and S1B). Through CAR T cell generation and flow cytometric grouping, we estimated that the CAR diversity was reduced from 10^6^ to 10^4^ (Figure 1A).

### Cytotoxicity-based pooled screening

The goal of pooled screening was not precise ranking, but rapid enrichment of CAR variants with both strong cytotoxicity and proliferative capacity. We sought to evaluate a large number of CARs using a pooled screening strategy; however, no established method existed for quantitatively measuring cytotoxicity at the single–cell level. To address this issue, we utilized the Beacon™ Optofluidic System (Bruker Cellular Analysis), which can isolate single live cells, perform real-time imaging, and unload selected cells. We first sought to detect CAR T cell–mediated cytotoxicity against target cancer cells at the single-cell level using the Beacon™ platform. Upon several contacts between CD19 CAR T cells and GFP-expressing Nalm6 cells cultured on Beacon, caspase-3-associated apoptosis was observed in Nalm6 cells within several hours, concomitant with the loss of GFP signal (Figure 2A). To enable simultaneous assessment of CAR T cell cytotoxicity and proliferation, we optimized the loading conditions for CD19 CAR T and Nalm6 cells (Figure 2B). Under these conditions, >50% and >80% of pens contained one and several CAR T cells, respectively, and the average number of Nalm6 cells per pen was 11.2. After a 3-day incubation, GFP signal in Nalm6 cells disappeared in pens loaded with CD19 CAR T cells, whereas Nalm6 cells continued to grow in pens not loaded with CD19 CAR T cells. Furthermore, CAR T cell proliferation, which was undetectable under short-term culture (< 1-day), became evident upon extended incubation (Figure 2C). Together, these results indicate that a single-cell assay on Beacon™ can quantitatively evaluate both the cytotoxicity and proliferative capacity of CAR T cells. Under optimized assay conditions, screening of CD19 CAR T variants generated by the MDPD method, with FACS-based fractionation into CD62L^high^ and CD62L ^low^ CAR T populations (Figure S1B), identified CAR T cells exhibiting strong cytotoxicity and high proliferative activity in both populations (Figure 2D). Screening hits based on cytotoxic (100%) and proliferative (>600%) activities were selected and then unloaded from the device. Through preliminary validation of filter conditions using a plasmid pooled coding known CAR genes (Figure S1C), we confirmed cutoff conditions that achieve both a high recovery rate and a low error rate for pools including 288 or fewer CARs. We unloaded 12 pools, with each pool containing CAR T cells from 6 to 25 pens for a total of 286 pens. We then extracted the mRNA from each pool and performed amplicon sequencing of the mutated region (Figure 2E). We selected 72 samples exhibiting high cytotoxic and proliferative activity from each of the CD62L^high^ population and the CD62L^low^ population. In total, 144 samples were selected in descending order based on the number of NGS reads in each pool.

**Figure 2:**
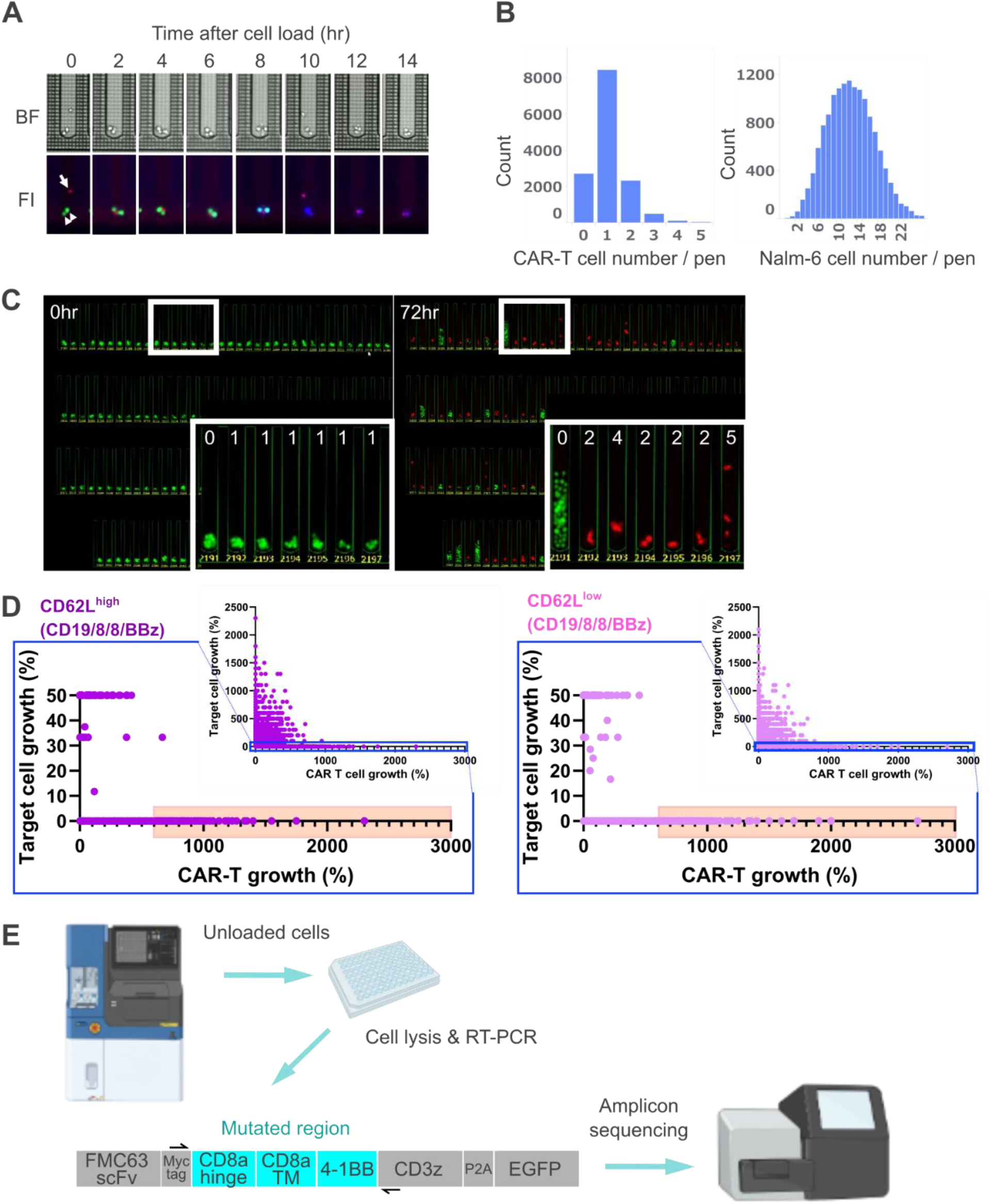
Cytotoxicity-based pooled screening. (A) Coculture of CAR T labeled with CellTrace Far Red and GFP-expressing Nalm6 in culture media including caspase indicator. Blue indicates caspase activation. The arrow and arrowhead indicate CAR T and Nalm6 cells, respectively. BF; Bright field image, FI; Fluorescence image. (B) Histograms of CAR T (left panel) and Nalm6 (right panel) cell number distribution right after cell loading. (C) Representative fluorescent images at 0-hr (left panel) and 72-hr (right) culture. Red and Green signals indicate CD8^+^ CAR T and Nalm6, respectively. An overview and magnification images of white area. Numbers in magnification image indicate the number of CAR T cells at 0 and 72 hr. (D) Scatter plots in CAR library screening for CAR T cell growth (X-axis) and cytotoxicity (Y-axis). Left panel and right panels show CD62L^high^ and CD62L^low^ populations, respectively. The large graphs magnify the high-cytotoxicity areas, which are indicated by the blue squares in each of the small graphs on upper right. “Target cell growth = 0%” indicates 100% cytotoxicity. Samples plotted in orange highlighted area (target cell growth = 0% and CAR T cell growth > 600%) were selected for unloading. (E) Schema of amplicon sequencing unloaded screening hits. Created with BioRender.com.

### Automated arrayed screening system

To further validate the cytotoxicity of CAR T cells identified through pooled screening, we developed an arrayed screening system that enables quantification of cytotoxicity using bulk CAR T cell populations, because single-cell phenotypes are heavily influenced by host cell heterogeneity. To obtain reliable and reproducible cytotoxicity data in a high-throughput manner, we established a laboratory automation system capable of performing the entire process of CAR T cell assay, from virus vector production to detection of target cell killing (Figure 3A). Our automation setup includes a liquid handler, CO2 incubators, a centrifuge, a flow cytometer, and a microplate reader, each coordinated by a single-arm plate handler and shuttles (Figure S2A). This laboratory automation system was used to produce viral vectors in quadruplicate and to transduce T cells derived from two independent donors with two virus replicates, respectively. In total, we obtained four CAR T cell replicates (Figure 3B). We measured cell concentration and the proportion of GFP-positive cells to determine transduction efficiency using the flow cytometer. Subsequently, the liquid handler diluted transduced T cells and mixed them with luciferase-expressing Nalm6 target cells in quadruplicate. Cytotoxicity was evaluated by measuring luminescence 23 hours after co-culture (1-day killing). To assess serial killing activity, the addition of target cells and luminescence measurement were repeated over a 10-day period (Figure 4B). This automated system can process 288 samples per cycle, excluding assay controls, and completes the entire process within approximately one month per screening cycle. Data stability and assay suitability are critical metrics in HTS. Most T cell transduction efficiencies observed during this screening were sufficient to validate CAR T cell function (Figure 3C). In the 1-day killing assay, the signal-to-background ratio (S/B) and a screening window coefficient, Z’- factor were consistently high across plates, indicating that this assay is well-suited for HTS (S/B > 10 and Z′ > 0.5) (Zhang et al., 2012) (Figure 3D & 3E).

**Figure 3:**
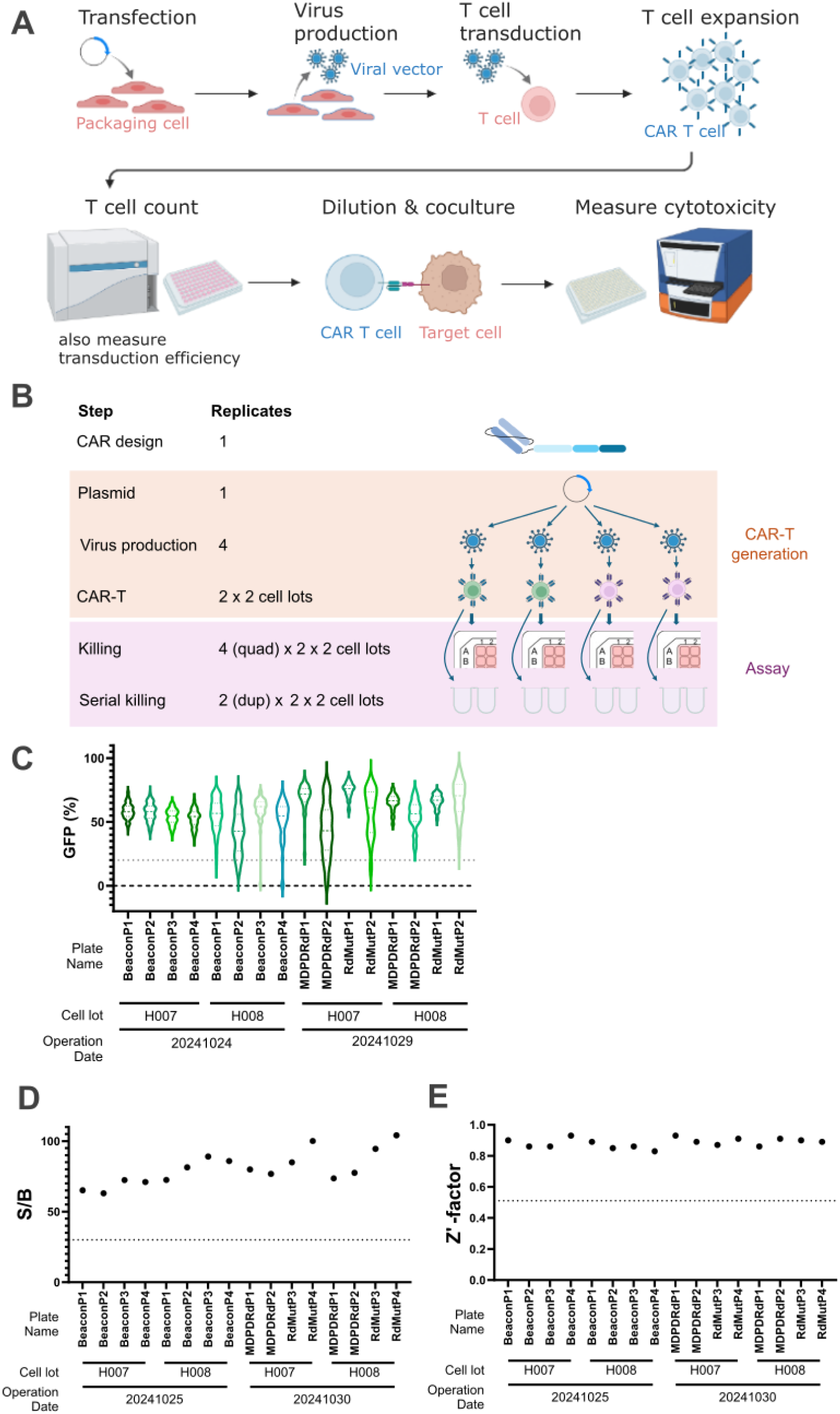
Automated cytotoxicity assay system for arrayed screening. (A) Schematics of whole process of automated CAR T production and evaluation. Created with BioRender.com. (B) Schematics of sample replication strategy in this screening. Created with BioRender.com. (C) Distribution of CAR T cell transduction efficiency across plates. The violin plot indicates distribution of GFP expression levels in each plate. (D) & (E) These dot plots show stability of 1-day killing data in each plate. Dot lines indicate internal criteria; over 0.75 of Z-factor (C) and over 30 of signal/noise ratio (S/B) (D). To calculate Z-factor and S/B, we set negative control of wells added Nalm6-Luc but no T cells, as well as positive control of wells with FMC63/28/28/28z CAR T cells and Nalm6-Luc at an E:T ratio of 2:1 in each plate.

**Figure 4:**
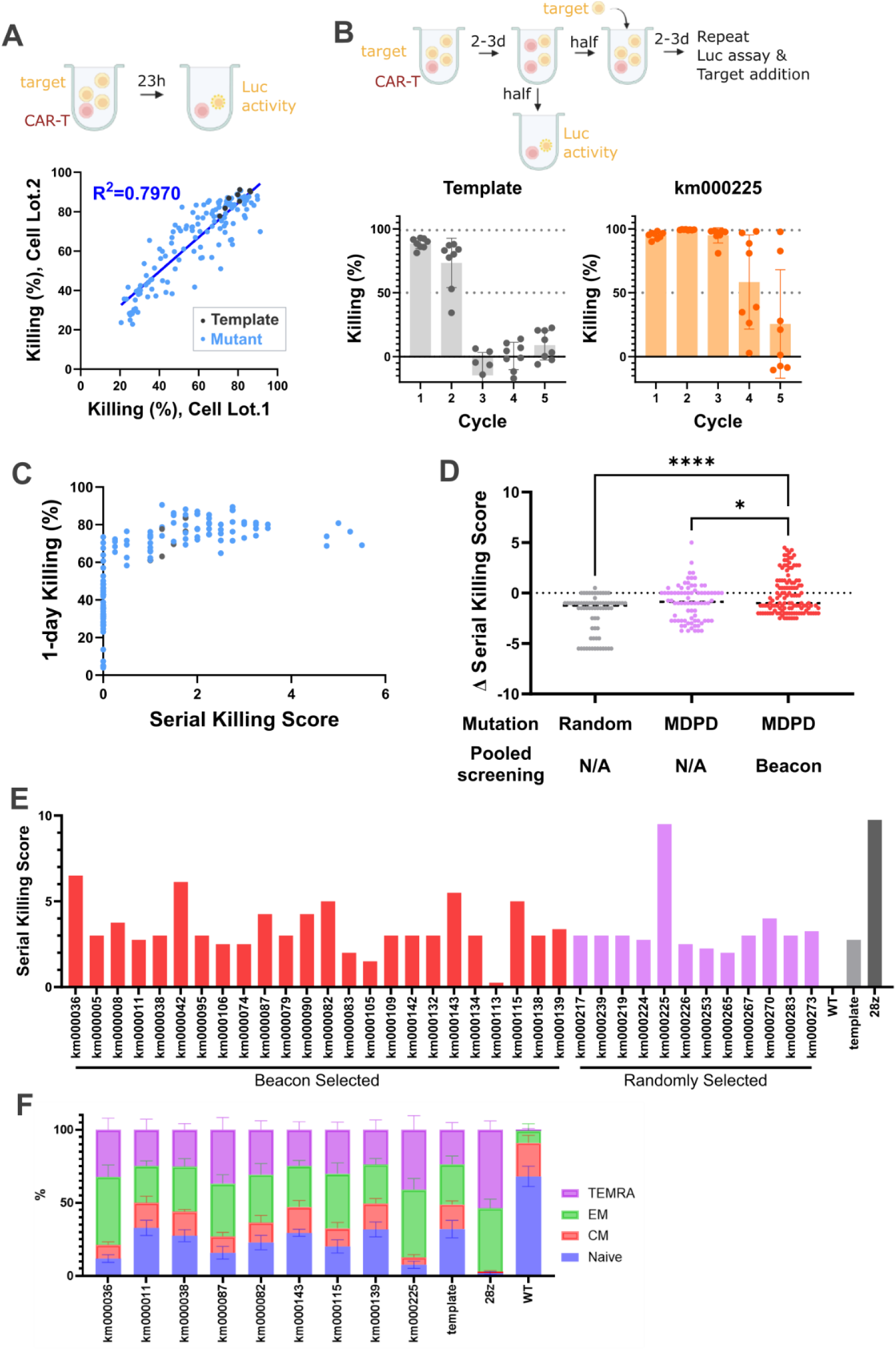
Serial killing assay for screening higher cytotoxic CARs. (A) Schematic of 1-day killing assay (upper). Created with BioRender.com. Scatter plot shows killing rate against target cell (Nalm6-Luc) in 1-day killing assay (E:T=1:8) and correlation the cytotoxicity across host T cell lots (lower). Blue dots indicate variants; gray dots indicate templates. The coefficient of determination (R²) derived from Pearson’s correlation is shown. (B) Schematic of serial killing assay (left). Created with BioRender.com. CAR T cells were cocultured with Nalm6-Luc at an E:T of 1:8 for 2-3 days, and half of coculture was reserved half volume for next cycle and remaining half is used to measure live target cells. This cycle was repeated 4 times. Bar graphs indicate killing rate at each cycle (right). Average±SD, N=8. (C) Scatter plot indicates correlation between 1-day killing rate and serial killing score. The serial killing score calculation formula is in the Method. (D) Dot plot indicates Δserial killing score of randomly mutated group (gray), MDPD-designed randomly selected group (pink), and MDPD-designed Beacon-selected group (red). The Δserial killing score calculation formula is in the Method. P values were determined using one-way ANOVA with Dunnet’s multiple comparison test. *, P=0.0129; ****, P<0.0001 (E) Bar graph indicates serial killing score of selected 36 variants and controls (gray). H001-009 were selected for *in vivo* experiments. WT, non-transduced T cell; 28z, FMC63/28/28/28z CAR T cell. (F) Stacked bar chart indicates the differentiation status of selected 9 variants by showing the populations of CD62L^high^/CD45RA^high^ (Naive), CD62L^high^/CD45^low^ (CM), CD62L^low^/CD45^low^ (EM), and CD62L^low^/CD45^high^ (TEMRA). Average±SD, N=4.

### Wide range quantitative cytotoxicity assay

Using the automated arrayed CAR T cell screening system, we performed cytotoxicity assay in which CAR T cells and luciferase-expressing target Nalm6 cells were cocultured for 23 hours, and then luminescence was measured to quantify live target cells. Only samples with a transformation efficiency of 20% or higher were included in downstream analyses, because killing activity decreased markedly below this threshold (Figure S3A). Killing rates measured using T cells derived from two independent donor were strongly correlated (Figure 4A). This assay demonstrated high resolution and reproducibility for lower-activity samples, revealing a loss-of-function mutation, K72E located in the membrane-proximal region of the 4-1BB intracellular domain (Figure S3B). However, high-activity CAR designs, including the template, exhibited signal saturation in the 1-day killing assay. To extend the quantitative dynamic range toward higher cytotoxic activity, serial killing assays were performed. In serial killing assay, CAR T cells were repeatedly challenged with luciferase-expressing Nalm6, and cytotoxic activity was assessed over multiple cycles (Figure 4B). Serial killing activity was quantified based the number of cycles showing ≥99% and ≥50% target cell killing (Figure 4C). Δserial killing score were calculated by subtracting the serial killing score of the template FMC63/8/8/BBz CAR from that of each sample within the same plate. Only CAR designs exhibiting top-tier activity in 1-day killing assay showed measurable serial killing activity and were clearly distinguished by the serial killing score. Based on Δserial killing scores, we compared randomly mutated CARs, MDPD-designed randomly selected CARs, and MDPD-designed Beacon™-selected CARs (Figure 4D). MDPD-designed randomly selected CARs showed significantly higher Δserial killing score than random mutated CARs, while Beacon™-selected CARs exhibited further enhancement. Beacon™-selected CARs subdivided based on CD62L expression during sorting; however, CD62L^high^ and CD62^low^ populations did not differ significantly in serial killing activity (Figure S3C). These results demonstrated that MDPD can effectively circumvent weak CAR designs, and Beacon™ selection can enrich CAR designs with high killing activity.

### Re-evaluation of selected CAR designs

To select CAR designs for *in vivo* validation, we confirmed cytotoxic activity of the 36 candidates identified by 1-day killing and serial killing data. We repeated serial killing assay on a single assay plate (Figure 4E), leading to selection of nine CAR designs; eight from Beacon™-selected group and one from MDPD-designed randomly selected group. Additionally, we also tested T cell differentiation status of each sample following coculture with CD19-expressing K562 cells (Figure 4F). All variants exhibited similar or reduced proportions of naïve and CM T cells compared to the template FMC63/8/8/BBz CAR. However, these variants retained higher levels than FMC63/28/28/28z CAR, which tends to differentiate quickly despite high cytotoxicity (Figure 4F and S4A). Given the known trade-off between CAR T cell memory phenotype and cytotoxicity (Cho et al., 2025), variants exhibiting strong cytotoxicity while maintaining moderate stemness were prioritized for further evaluation.

### *In vivo* evaluation

To validate our screening hits *in vivo*, we treated leukemic mice with the selected CAR T cells. Luciferase-expressing Nalm6 cells were intravenously injected into immunocompromised mice, and tumor growth was monitored for 35 days using bioluminescence imaging (Figure 5A). To ensure consistency with *in vitro* screening conditions, CAR T cells were generated from CD8^+^ T cells (Figure 5B and 5C) in addition to PBMC-derived CAR T cells (Figure S5A). Compared to the template FMC63/8/8/BBz CAR T cell, both the km000143 and km000225 CAR T cells exhibited reduced tumor burden and prolonged survival. Identification of these two variants with superior *in vivo* efficacy demonstrates the reliability of the cytotoxicity-based screening platform combined with AI-assisted CAR design.

**Figure 5:**
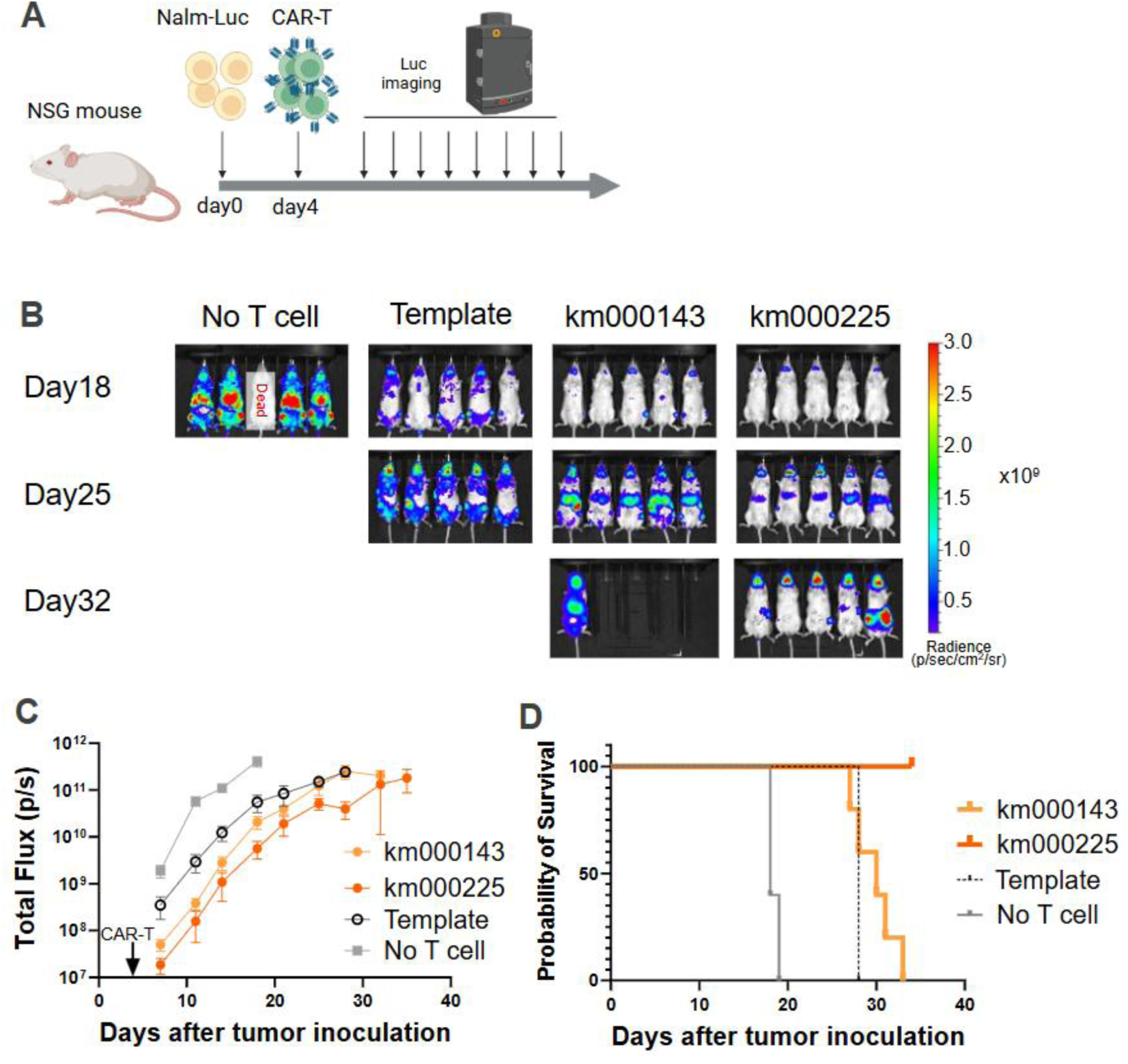
km000143 and km000225 CAR T cells exhibited enhanced antitumor activity against B cell lymphoma model. (A) Schematic design of leukemia model experiment. Created with BioRender.com. (B) Measurement of tumor progression using bioluminescence at indicated days after *i.v.* injection of 1 ×10^6^ Nalm-Luc cells. Mice were treated with PBS or 2 ×10^6^ CD8^+^ T-derived CAR T cells. (C) Tumor burden (total flux) quantified by photons/sec in mice treated with PBS or CD8^+^ T-derived CAR T cells. (D) Kaplan-Meier plot indicating overall survival of mice after tumor injection.

### Identification of essential mutations for enhancement

The km000225 CAR harbors eight mutations (A12V/I15P/S17L/L20A/H36N/L73I/I76L/M82L; Figure 6A), making it unclear which mutation(s) contributes to the functional improvement. To address this, we generated 144 CAR variants composing subsets of km000225-derived mutations and tested their cytotoxicity using serial killing assay. Most km000225-derived variants showed improved cytotoxicity compared with the template (Figure 6B and S6A). Notably, CARs containing both I15P and L20A mutations within the CD8α hinge domain (I15P/L20A) demonstrated markedly enhanced cytotoxicity relative to other variants (Figure 6B and S6B). A CAR containing only three mutations (I15P, L20A and H36N; km000491) exhibited cytotoxic activity comparable to km000225 (Figure 6B). We also verified T-cell stemness of these CAR variants, showing a graded trade-off relationship between cytotoxicity score and rate of CD62L^high^ population (Figure 6C). For *in vivo* validation, we selected km000527 harboring I15P, L20A, H36N and L73I based on its higher cytotoxicity and comparable stemness to the template FMC63/8/8/BBz CAR. In a xenograft model, km000527 demonstrated superior antitumor efficacy compared with FMC63/8/8/BBz CAR T cells (Figure 6D & 6E). Altogether, these results demonstrated that the screening platform enabled identification of new CAR variants with enhanced *in vivo* efficacy from theoretically 10^8^ variants. Furthermore, it facilitated the systematic identification of essential mutation combinations underlying functional enhancement.

**Figure 6:**
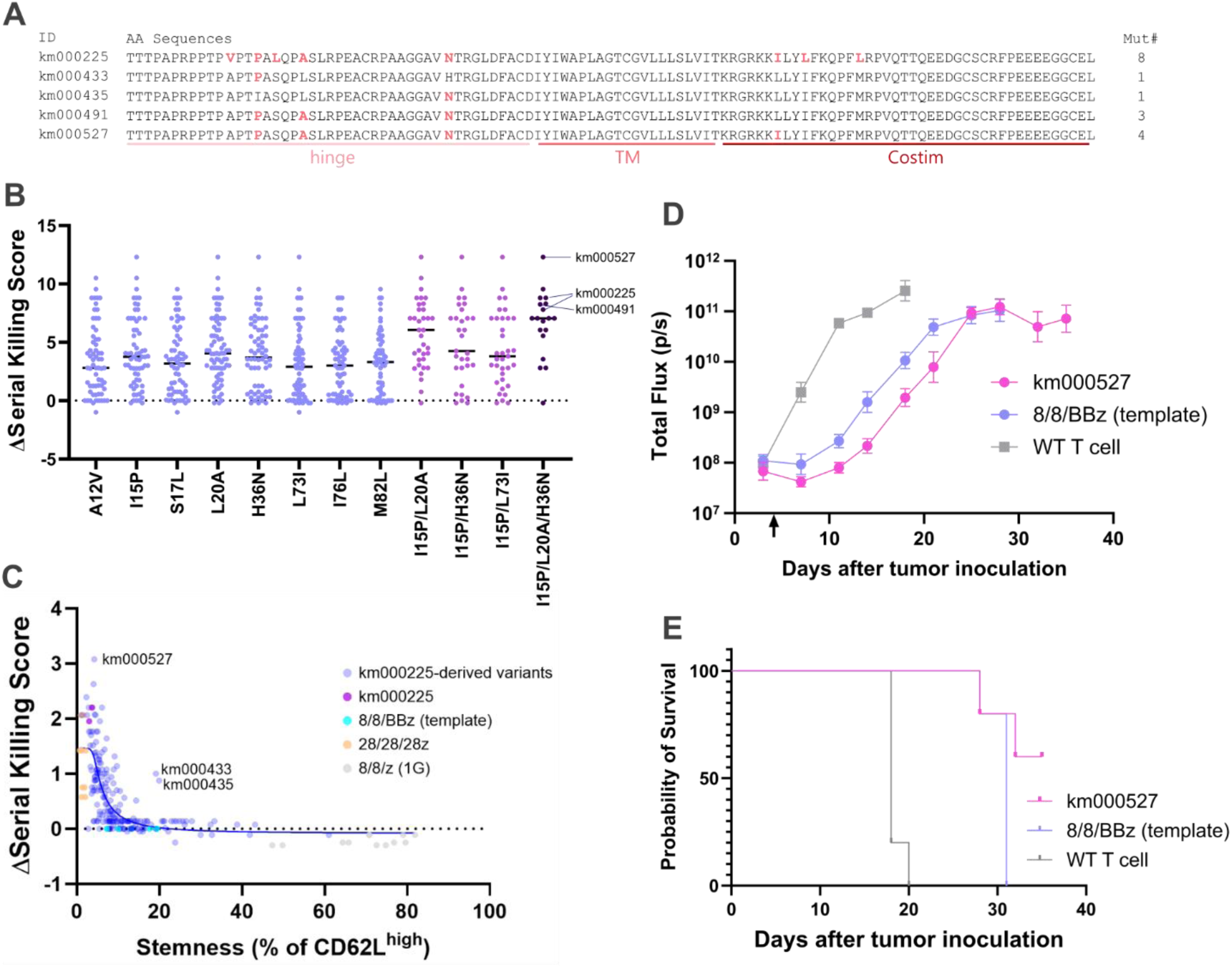
Essential mutations for improved activity and phenotypical difference. (A) Amino acid sequences of the various regions in the km000225 and km000225-derived variants mentioned in Fig.6. Red bold characters indicate mutations. (B) Scatter plot indicates Δserial killing score of km000225-derived variants. A12V denotes the group of variants containing A12V mutation. I15P/L20A denotes the group of variants containing both I15P and L20A mutations. (C) Dot plot indicates the correlation between Δserial killing score and he percentage of CD62L^high^ cells after 48-hour coculture with K562-CD19. The blue solid line indicated a curve fitted using a five-parameter logistic model. (D) Tumor burden (total flux) quantified by photons/sec in mice treated with non-transduced T cells (WT T cell) or PBMC-derived CAR T cells. (E) Kaplan-Meyer plot indicating overall survival of mice after tumor injection

## DISCUSSION

A systematic understanding of the CAR sequence–function relationship requires screening strategies that can simultaneously handle vast design diversity and provide quantitative functional resolution. In this study, we present not merely a new assay or a new CAR variant, but an integrated screening platform that combines *in silico* CAR design, cytotoxicity-based pooled screening, and automated, high-resolution arrayed evaluation. To validate our platform, we generated approximately 10⁸ variants based on the FDA-approved FMC63/8/8/BBz architecture using MDPD. We then screened a subset of these variants using our pooled and arrayed screening systems. Through cytotoxicity assays, we step-wisely narrowed the diversity of the CAR T cell library and selected nine variants for *in vivo* evaluation. Two variants out of these demonstrated superior *in vivo* efficacy compared to the template CAR. Additionally, our platform enabled the identification of an essential mutation combination underlying the enhanced performance observed in the top variant.

One of the methods frequently used to construct of new CAR libraries is domain shuffling, which may generate CARs with novel functions (Bloemberg et al., 2020; Castellanos-Rueda et al., 2025, 2022; Duong et al., 2013; Goodman et al., 2022; Gordon et al., 2025, 2022; Perez et al., 2025; Rios et al., 2023). However, as long as evaluation systems are built based on conventional readouts, it remains less likely to discover entirely new functions. Therefore, we used a mutation library derived from an FDA-approved CAR design and adopted a screening strategy focused on fine-tuning rather than discovering new functions. The primary challenge in creating such a mutation library is narrowing down the vast number of possible variants *in silico*, because testing all variants is impractical. By leveraging protein fitness prediction to guide mutation selection, our MDPD efficiently enriched functional variants while preserving a large theoretical design space.

In this study, we utilized cytotoxicity as a key indicator for screening CAR T cell function. While cytotoxicity is a crucial effector mechanism of CAR T cells, its low throughput and poorly defined quantitative range have limited its use as a primary screening metric. To address this, we developed a single-cell cytotoxicity pooled screening method. Beacon™ system enabled long-term cell monitoring, allowing us to quantify the number of target cells each CAR T cell has killed and to assess CAR T cell proliferation over time. This method, unlike other short-term qualitative single-cell cytotoxicity assays (Subedi et al., 2021; Wei et al., 2025), is well-suited for selecting CAR T cells with higher activity. In addition, we established an arrayed cytotoxicity screening method suitable for evaluating CAR designs with higher cytotoxic activity. Coupled with automation, this system not only increased throughput but also produced highly reproducible data for these complex, time-consuming experiments.

Previous studies have shown that enhanced short-term activity, including strong *in vitro* cytotoxicity, does not necessarily lead to improved efficacy, as excessive CAR signaling can induce T cell exhaustion and impair long-term persistence (Guedan et al., 2020; Long et al., 2015; Yin et al., 2023). Therefore, we integrated T-cell stemness evaluation into the arrayed screening (Fig.6D). Various types of combinations of cell surface markers (e.g. CD62L and CD45RA) serve as indicators of T-cell stemness/differentiation status. Many studies have shown that CAR T cell differentiation status correlates with CAR T cell persistence *in vivo* (Drent et al., 2019; Long et al., 2015; Sommermeyer et al., 2016; Xu et al., 2014), and in some cases, with clinical outcomes (Biasco et al., 2021; Fraietta et al., 2018; Kasuya et al., 2024; Raje et al., 2021). Interestingly, it was widely known in immunotherapy field there is a trade-off relationship in T cells between short-term activity (e.g. cytotoxicity) and long-term activity (e.g. stemness) (Cho et al., 2025; Conti et al., 2026; Kang et al., 2024; Rodriguez et al., 2026). However, this study demonstrates the graded trade-off relationship based on variations in CAR design. The screening strategy, which assesses both cytotoxicity and stemness, is well-suited for tailoring CAR functions and also identifying CARs that exhibit exceptional high performance in both parameters simultaneously.

Overall, our screening platform, which utilizes *in silico* design and HTS, enabled the identification of CAR variants that exhibit functional advantages. We will combine the screening platform with computational frameworks for predicting CAR functions to run further optimization cycles (Yoshida et al., 2026). The platform can be applied to comprehensively identify functionally essential motifs within existing CARs (Figure.S3B) and mutations critical for functional enhancement in new CAR variants (Figure.6). Furthermore, it paves the way for advancing our understanding of CAR design principles and for maximizing the intellectual property related to CAR sequences.

## MATERIALS AND METHODS

### CAR design strategy

EVcouplings(Hopf et al., 2019) was used to generate multiple sequence alignments (MSAs) and to train EVmutation models for fitness prediction. The amino acid sequences of the CD8α hinge and the 4-1BB costimulatory domain were analyzed separately. For each domain, MSAs were built against the UniRef100 database using EVcouplings, with bitscore inclusion thresholds of 0.4 (CD8α hinge) and 0.3 (4-1BB costimulatory domain). Using EVmutation models trained on these MSAs, we computed predicted fitness scores for all single-residue substitutions in each domain. For each domain, we selected higher-fitness substitutions such that the combined design space comprised up to 10^8^ substitution combinations. In total, this yielded 73,903,104 designed variants of FMC63/8/8/BBz. The resulting designs were submitted to Twist Bioscience as a Combinatorial Variant Library for synthesis.

### Pooled vector construction

A mutation dsDNA library from CD8α hinge and transmembrane and 4-1BB costimulatory domain was synthesized by Twist Biosciences. A pHR lentiviral vector containing an SFFV promoter followed by FCM63, BamHI site, CD3ζ intracellular domain, P2A ribosomal skipping site, and EGFP was used as a backbone. The mutation dsDNA library was subcloned to the BamHI digested backbone using NEBuilder HiFi DNA Assembly Master Mix (NEB) according to manufacturer’s protocol. The cloning product was transformed into HST08 competent *E. coli* cells (Takara). Transformed cells were plated LB agar plate (Nacalai tesque) containing carbenicillin (FUJIFILM Wako Chemicals), over 20 million colonies were scraped, and the plasmids were extracted using NucleoBond Xtra Midi kit (Takara).

### Cell lines

Nalm6 cell was obtained from RIKEN BioResource Center (BRC) and stably transduced with EGFP and firefly luciferase using the lentiviral vector pHR-SFFV-Luc-p2A-EGFP. K562 cell was also obtained from RIKEN BRC and stably transduced with truncated CD19 using the lentiviral vector pHR-SFFV-tCD19. Nalm6 and K562 were cultured with RPMI1640 (Nacalai tesque) supplemented with 5% FBS (Corning), 2 mmol/mL L-glutamine, 100 U/mL penicillin and 100 µg/mL streptomycin (Nacalai tesque). 293FT cells (ThermoFisher) were cultured in DMEM (Nacalai tesque) supplemented with 10% FBS, L-glutamine, 1 mmol/mL sodium pyruvate, 100 U/mL penicillin and 100 µg/mL streptomycin (Nacalai tesque).

### Primary T cell isolation and culture

Human PBMCs were obtained from STEMCELL technologies. CD8^+^ T cells were obtained from STEMCELL technologies and were isolated from LeukoPack (STEMCELL technologies) with EasySep RBC depletion reagent and EasySep Human CD8+ T Cell Isolation Kit (STEMCELL technologies).

PBMCs and CD8+ T cells were maintained in cX-VIVO, X-VIVO15 (Lonza) supplemented with 5% human AB serum (Millipore-Sigma), 100 μM 2-mercaptoethanol (FUJIFILM Wako Chemicals), 25 U/mL IL-2 (Peprotech), 10 ng/mL IL-7, and 10 ng/mL IL-15 (Miltenyi biotec).

### Lentiviral transduction of T cells

293FT cells were suspended with complete X-VIVO15 supplemented with 5% human AB serum and 100 µM 2-mercaptoethanol and transfected with pHR-GOI vector and packaging plasmid mix; pMD2.G encoding for VSV-G pseudotyping coat protein, pCMVR8.74, and pAdvantage (Promega), using Lipofectamine 3000 (ThermoFisher) or FuGENE 4K (Promega). Virus supernatants were harvested 48 hours after the transfection. CD8^+^ T cells were activated using Dynabeads Human T-cell Activator CD3/CD28 (ThermoFisher) at 25 µL per one millinon T cells. Twenty-four hours after activation of T cells, the virus supernatant was added to the activated T cells. Dynabeads were removed 2 days later. Transduction efficiency was evaluated 3 days after removal of Dynabeads and coculture with target cells were performed the following day.

### Cell sorting

The CAR T cell pool was co-cultured with CD19-expressing K562 cells at a 1:1 effector-to-target (E:T) ratio for 48 hours. After coculture, cells were stained with APC conjugated anti-CD45RA (Biolegend, 304150), BV421 conjugated CD62L (Biolegend, 304827), PE conjugated CD8 (Biolegend, 344706), and BV785 conjugated CD39 (Biolegend, 328240) antibodies. CAR expression was detected by EGFP signal. Cell sorting was performed using the FACSMelody (BD bioscience), and the data were analyzed FlowJo version10 (BD bioscience).

### Beacon™ screening

CAR T cells and Nalm6 cells, prepared at 4 × 10^6^ cells/mL and 2 × 10^7^ cells/mL, respectively, were loaded onto the Beacon™ OptoSelect 3500 chip (Bruker Cellular Analysis). During on-chip assessment, cells were cultured in cX-VIVO. For loading, cX-VIVO containing 30% Loading Reagent (Bruker Cellular Analysis), which mitigates light-induced cytotoxicity was used. Apoptosis was monitored using the caspase indicator, NucView 405 (Biotium) at a final concentration of 2 µM in cX-VIVO. For real-time imaging of CD19 CAR T cells and GFP-expressing Nalm6 cells in the presence of the caspase indicator, CAR T cells were pre-stained with CellTrace Far Red (Invitrogen) at a final concentration of 500 nM. In pooled variant library screening, after 72 hours of co-culture, CAR T cells were stained with PE-conjugated CD8 antibody (BioLegend). The number of CAR T and Nalm6 cells was quantified by PE and GFP signals, respectively. CAR T cell proliferation was calculated from CAR T cell counts at 0 and 72 hours of culture, and cytotoxicity was calculated from Nalm6 cell counts at the same time points.

### Amplicon sequencing

The pool of hit cells evaluated by Beacon™ was unloaded into a 96-well plate containing lysis buffer and extracted the RNA. RT reaction and cDNA pre-amplification were performed using the SuperScript IV Single Cell/Low Input cDNA PreAmp Kit (Invitrogen) according to the manufacturer’s protocol. cDNA pre-amplification was performed using primer (forward, 5’-gtggtagctatgctatggac-3’; and reverse, 5’-gctcgttatagagctggttc-3’) with TruSeq Universal adapter (Illumina) for 18 cycles at the following conditions: denaturation 98°C for 30 sec, then cycling 98°C for 10 sec, 65°C for 10 sec, 67°C for 3 min, and a final extension for 5 min. High-fidelity PCR amplification using the adapter sequences was then performed using Platinum SuperFi II DNA Polymerase (Invitrogen) for 25 cycles at the following conditions: denaturation 95°C for 30 sec, then cycling 98°C for 10 sec, 60°C for 10 sec, 72°C for 15 sec, and a final extension for 60 sec. The concentration and size of the cDNA were quantified using a Bioabalyzer (Agilent). The amplified cDNA was sequenced using MiSeq (illumina). Sequencing was performed using 300-bp paired-end sequencing by Bioengineering Lab. Co., Ltd. A contig was created from the overlapping sequences of the paired reads to determine the full-length sequence of the analyzed region. Filters were set to remove sequence information resulting from PCR errors and sequence reading errors from the obtained sequence data. The filter criteria were 100% homology in unmutated regions, the absence of nonsense mutations in the amplified region, the absence of in-flame or flame-shift mutations in the amplified region, the presence of the resulting amino acid conversion sequence in the reference sequence, and the number of reads for each sequence being equal to or greater than the pre-validated cutoff (Figure S1C).

### Arrayed vector construction

Selected variants were re-synthesized using eBlocks (Integrated DNA Technologies). The dsDNAs were individually subcloned to the BamHI digested backbone (FMC63-(BamHI)-CD3ζ-P2A-EGFP) with NEBuilder HiFi DNA Assembly Master Mix. We transformed the cloning product into HST08 competent E. coli cells (Takara). Transformed cells were plated LB agar plate with carbenicillin, each single colony picked up, and each plasmid extracted with QIAprep Spin Miniprep Kit by QIAcube connect (Qiagen). Sequences of cloned fragments were confirmed by sangar sequencing (performed by Eurofins genomics).

### Arrayed screening with liquid handler

All plasmids were diluted to 100 ng/µL with nuclease-free water by pipetting robot Andrew+ (Andrew Alliance). 293FT cell suspension, diluted transfection reagent and diluted packaging mix were prepared by manual operation. 293FT cells were aliquoted to 96-well plate (IWAKI), and each plasmid was transfected to 293FT cells by liquid handler Biomek i7 (Beckman coulter). Frozen CD8+ T cells were thawed and activated for 24 hours with Dynabeads by manual operation. Virus supernatants from transfected 293FT cells were collected and mixed with activated CD8+ T cells by Biomek i7. Two days later, Dynabeads were removed by Biomek i7. cX-VIVO were added to T cells 6 days after T cell stimulation, and then, to measure cell concentration and transduction efficiency, flow cytometry analysis (CyteFLEX S) was performed overnight.

Ten µL of 1 ×10^6^ cells/mL luciferase-expressing Nalm6 (Nalm-Luc) were aliquoted to white 384-well round bottom plate (Sumitomo Bakelite). Transduced T cells were counted by the flow cytometer. T cell concentration was adjusted 1.25 ×10^5^ and 6.25 ×10^4^ cells/mL in cX-VIVO, and then the diluted T cells were added 10 µL to each well containing Nalm-Luc. 23 hours after starting coculture, 10 µL of OneGlo (Promega) were added to each well, and luminescence was measured with SpectraMax i3x (Molecular Device).

Fifty µL of 5 ×10^5^ cells/mL Nalm-Luc were aliquoted to 96-well round bottom plate (VIOLAMO). T cell concentration was adjusted 6.25 ×10^4^ cells/mL in cX-VIVO, and then the diluted T cells were added 50 µL to each well containing Nalm-Luc. Every 48 or 72 hours, 50 µL of cultured cell suspension was transferred to new 96-well round bottom plate and mixed with 50 µL of 5 ×10^5^ cells/mL Nalm-Luc. 20 µL of left cell suspension was transferred to black 384-well plate. Ten µL of OneGlo were added to each well, and luminescence was measured with SpectraMax i3x (Molecular Device). Serial killing score and Δserial killing score was determined using the following formula:

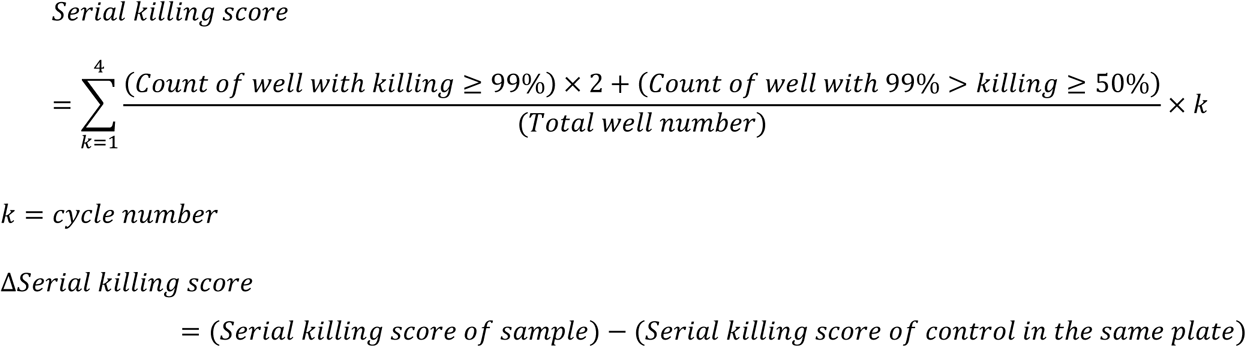

Metrics used to assess data quality included the Z’-factor, the signal-to-background ratio (S/B), and the coefficient of variation (CV). The Z′ (Z’-factor), S/B, and %CV values were calculated using the following equations, respectively.

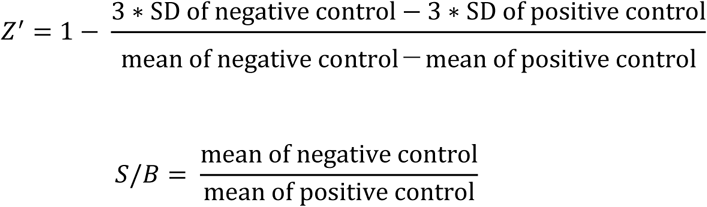

Fifty µL of 5 ×10^5^ cells/mL CD19-expressing K562 (K562-CD19) were aliquoted to 96-well round bottom plate. T cell concentration was adjusted 1.25 ×10^5^ cells/mL in cX-VIVO, and then the diluted T cells were added 50 µL to each well containing K562-CD19. Forty-eight hours after starting coculture, cells were stained with APC conjugated anti-CD45RA, BV421 conjugated CD62L, PE conjugated CD8 antibodies using microplate centrifuge (Agilent), plate washer (Biotek 405TS, Agilent), and liquid handler, and then were analyzed by flow cytometry (CytoFLEX S, Beckman Coulter) sequentially.

### *In vivo* tumor model

NOD.Cg-*Prkdc*^scid^ Il2rg^tm1Wjl^/SzJ (NSG) female mice (Jackson laboratory Japan) at 6 weeks old were intravenously (i.v.) injected with 1 million firefly luciferase-expressing Nalm6 cells. 3 days after tumor transplantation, the mice were grouped according to their body weight using EXSUS 10.1 statical analysis system, one million PBMC-derived or two million CD8^+^ T cell-derived CAR T cells were i.v. administrated on the next day. For monitoring tumor burden, VivoGlo Luciferin, In vivo Grade (Promega, P1043) was dissolved at 15 mg/mL in PBS and injected intraperitoneally at a concentration of 10 mg/kg based on body weight. Mice were anesthetized with isoflurane, and bioluminescence imaging was performed 15 min after injection in the IVIS Lumina II (PerkinElmer). Bioluminescence was quantified using Living Image Analysis Software v.4.7.3 (PerkinElmer). Mice were euthanized when they display deterioration of systemic condition presumably induced by high tumor burden or >20% weight loss. All animal experiments were conducted by Axcelead Drug Discovery Parteners in compliance with Institutional Animal Care and Use Committee (IACUC) in the institute.

## DATA AVAILABILITY

The datasets generated and/or analyzed during the current study are available from the corresponding author upon reasonable request. Access may be restricted due to confidentiality, intellectual property, or third-party licensing obligations. Where appropriate, data will be shared under a data use agreement.

## ACKNOWLEDGEMENTS

The authors are grateful to Hiroko Hanzawa, and Tomohiko Okuda for their leadership and managerial support, which enabled the successful completion of this study. We thank Axcelead Drug Discovery Parteners for performing the animal experiments.

## AUTHOR CONTRIBUTIONS

A.O. conceived the entire screening flow. S.H. designed the *in silico* CAR design strategy. S.M. and A.O. designed highly diverse plasmid pool construction protocol and high-throughput plasmid construction system. S.M. performed plasmid constructions and prepared CAR T cells for pooled screening and *in vivo* experiments. A.O. designed sorting protocol as the prescreening. Y.I. designed and performed pooled screening using Beacon™. T.K. designed and performed NGS analysis. T.Y. and A.O. built the laboratory automation system and prepared the protocols for arrayed screening. Y.N. performed experiments for NGS and operated the laboratory automation system. T.I. performed experiments using cell sorter and Beacon™. D.I. managed and analyzed the *in vivo* experiments. A.O. performed the manual evaluation of CAR T cells after screening. K.Y. and A.O. analyzed the data. S.T. supervised the research. A.O. created figures and wrote the manuscript. All authors commented on and approved the manuscript.

---

**Figure S1:**
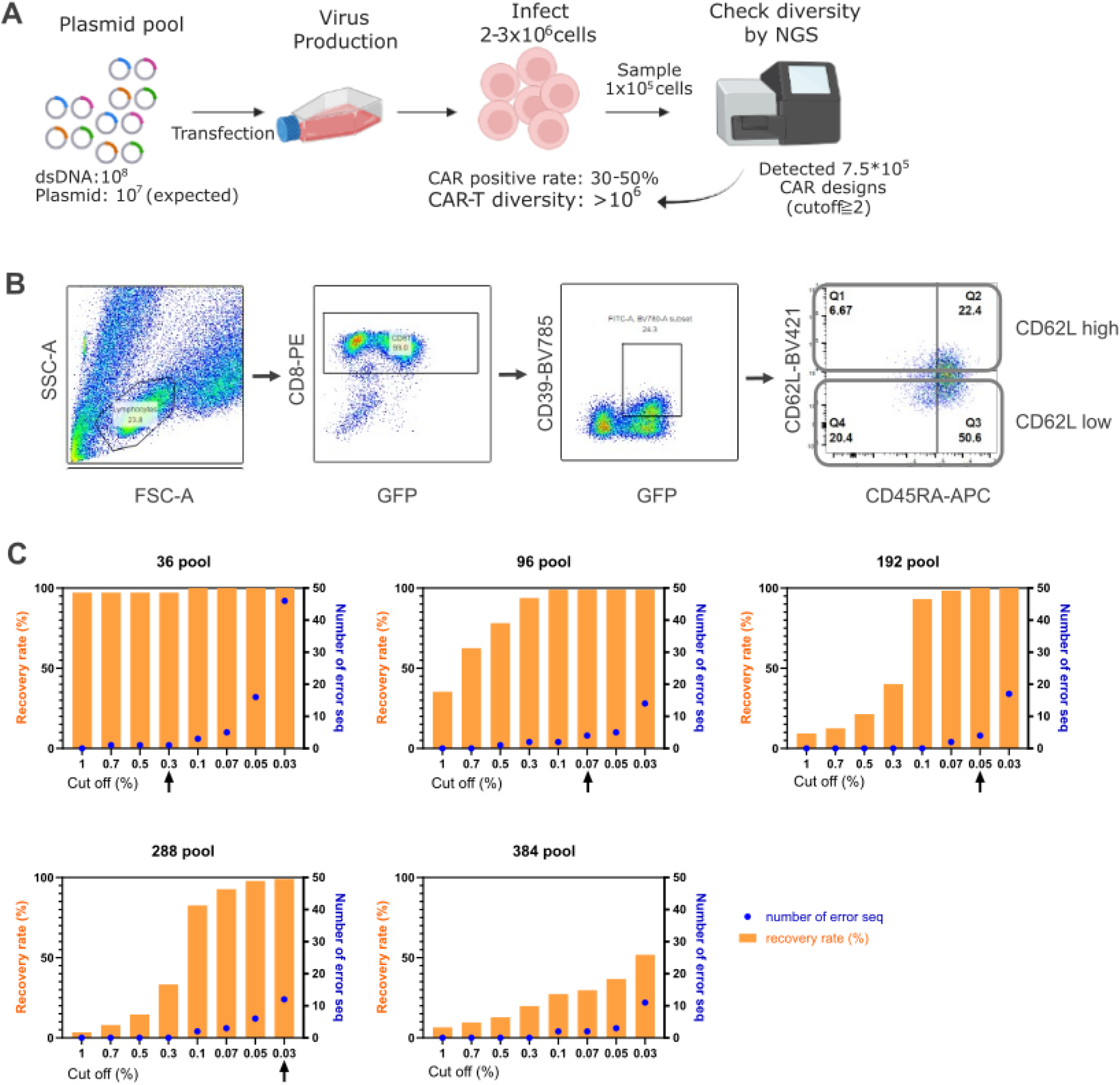
Confirmation of the CAR diversity, prescreening strategy, and optimization of NGS setting for pooled cell library. (A) Schematic of sampling procedure for diversity verification of the CAR T cell pool. Created with BioRender.com. (B) Flow cytometry plots show gating strategy for sorting before loading into Beacon. CAR(GFP)^+^/CD8^+^/CD39^+^/CD62L^high^ population and CAR(GFP)^+^/CD8^+^/CD39^+^/CD62^low^ population were sorted. (C) Bar graphs and dots graphs indicate NGS recovery rate and the number of error sequences of each pool, respectively. We examined filter conditions using pooled cell library transfected with known CAR genes. In each pooled cell library, we observed a tendency for the number of error sequences to increase exponentially as the cutoff was lowered. Furthermore, we observed a tendency for the recovery rate to plateau above 90%. Therefore, we considered the optimal cutoff for each cell population to be a cutoff before the number of error sequences increases exponentially and at a recovery rate of 90% or higher. Therefore, we considered appropriate cutoff conditions to be 0.3% for the NGS library pool including less than 66 CARs, 0.07% for the 67-144 pool, 0.05% for the 145-240 pool, and 0.03% for the 241-300 pool. On the other hand, in the 384 pool, even at the cutoff condition where error sequences begin to increase exponentially, the recovery rate was approximately 50%, suggesting that lowering the cutoff further could result in the detection of a large number of error sequences. Therefore, in this study, we set a 300 pool as the upper limit for gene sequence identification per well.

**Figure S2:**
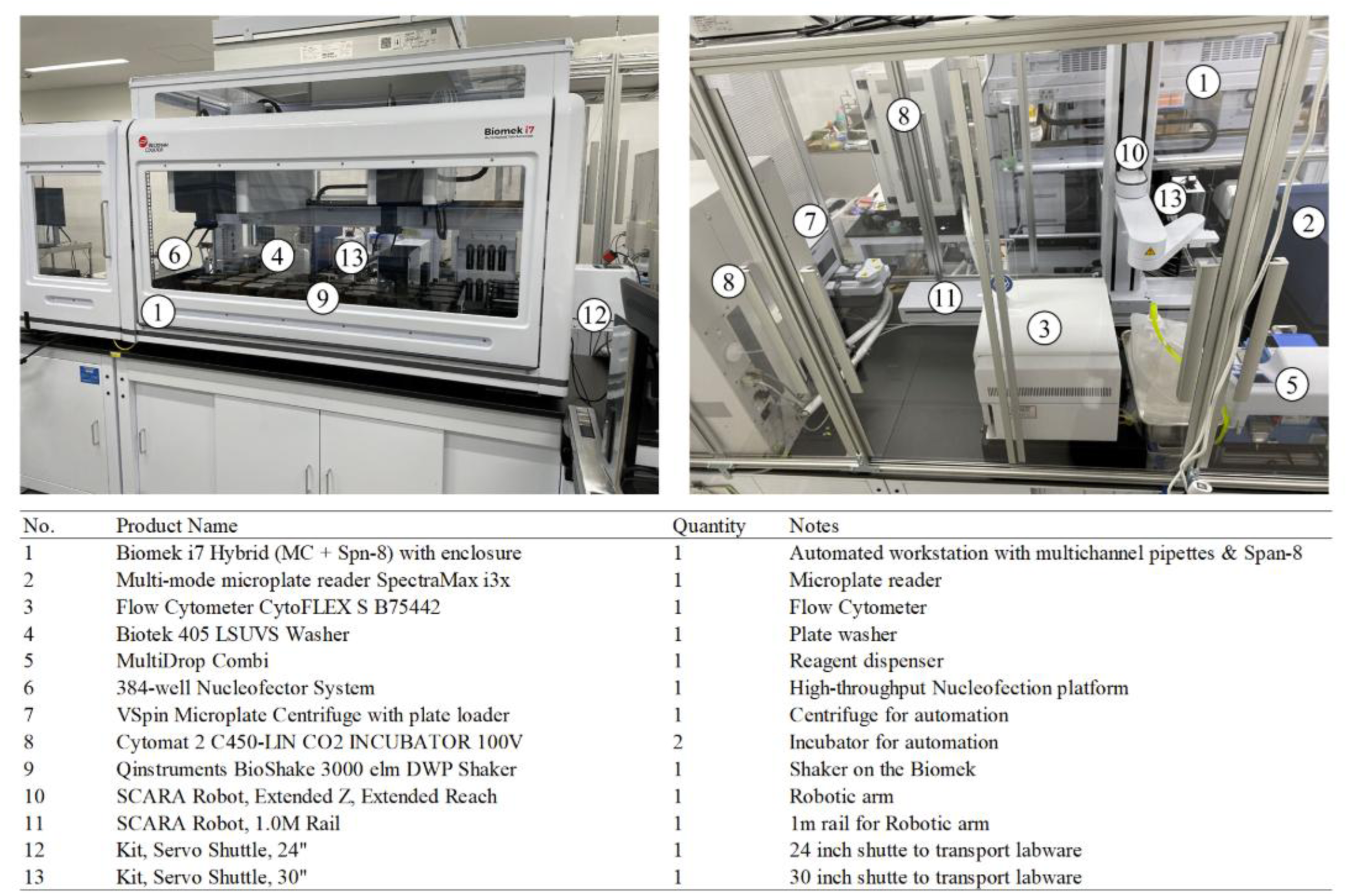
Hardware configuration of the arrayed screening system. Top: External view of the arrayed screening system. Bottom: List of instruments integrated into the arrayed screening system.

**Figure S3:**
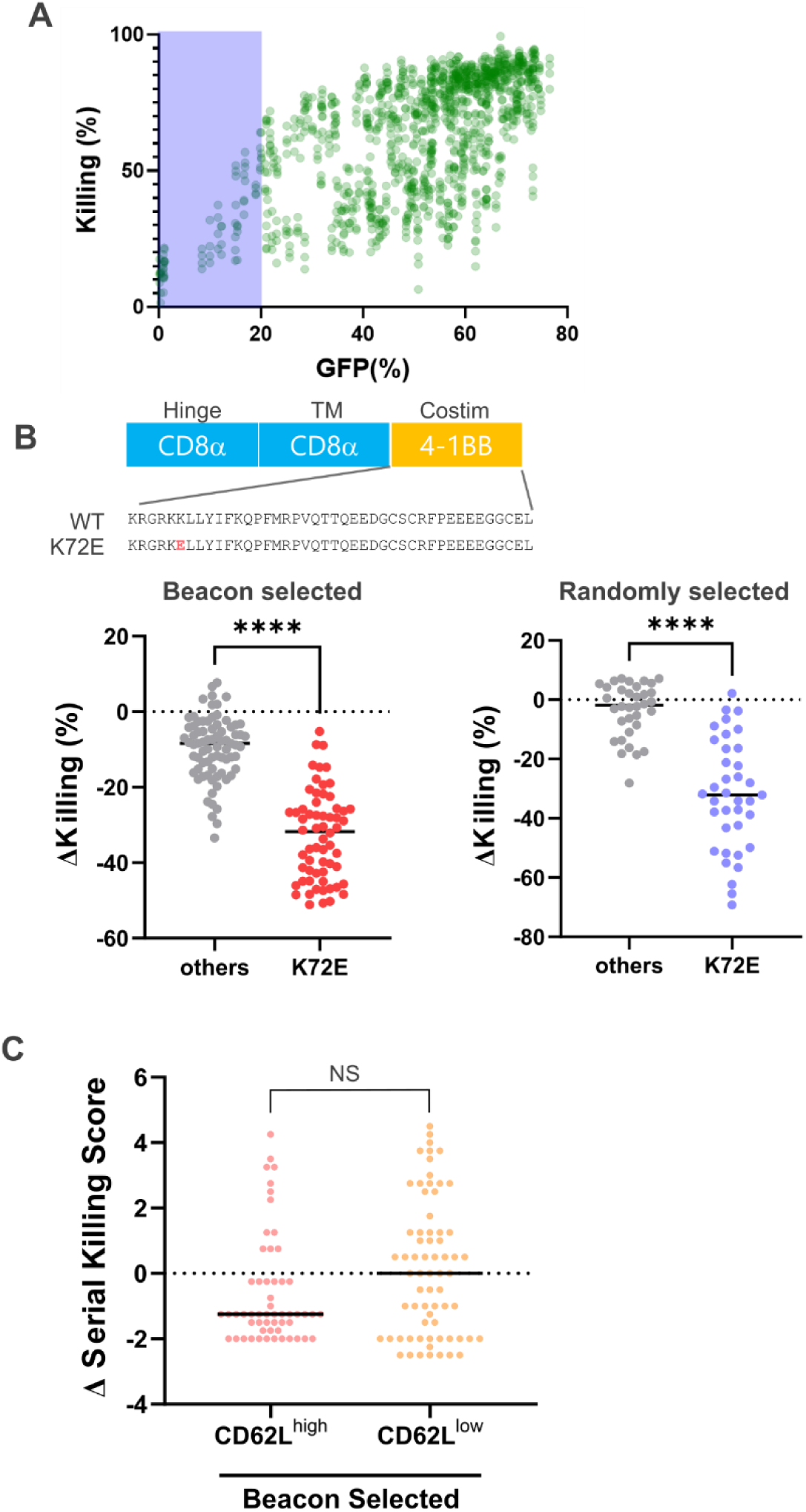
Trends revealed by cytotoxicity-based arrayed screening. (A) Scatter plot shows correlation between transduction efficiency (GFP (%)) and 1-day killing activity (killing (%)) of all samples in this screening including controls. (B) Upper schematic shows K72E mutation site in the 4-1BB–derived intracellular domain. Dot plots show Δkilling value of variants. Δkilling value was compared between variants with and without K72E among beacon selected group (left) and randomly selected group (right). P values were determined using unpaired t test with Welch’s correction. ****, P<0.0001 (C) Dot plot shows Δserial killing score for CD62L^high^ group and CD62L^low^ group which were grouped in the sorting step (Figure S1B). In Figure 4D, beacon selected group includes both groups CD62L^high^ and CD62L^low^ in this graph. NS, not significant.

**Figure S4:**
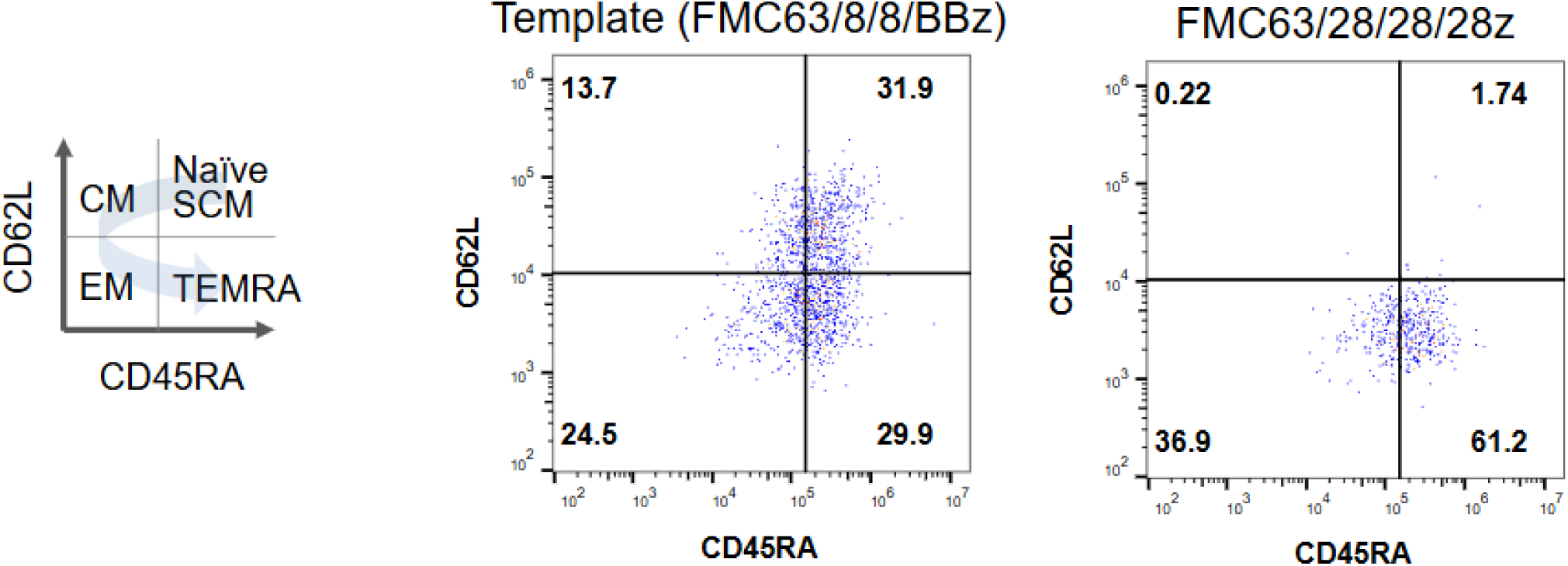
The difference of stemness between CAR designs. Representative flow cytometry plots showing CD45RA and CD62L expression in Figure 4F. Left diagram visualizes which differentiation status are plotted where in CD45RA/CD62L.

**Figure S5:**
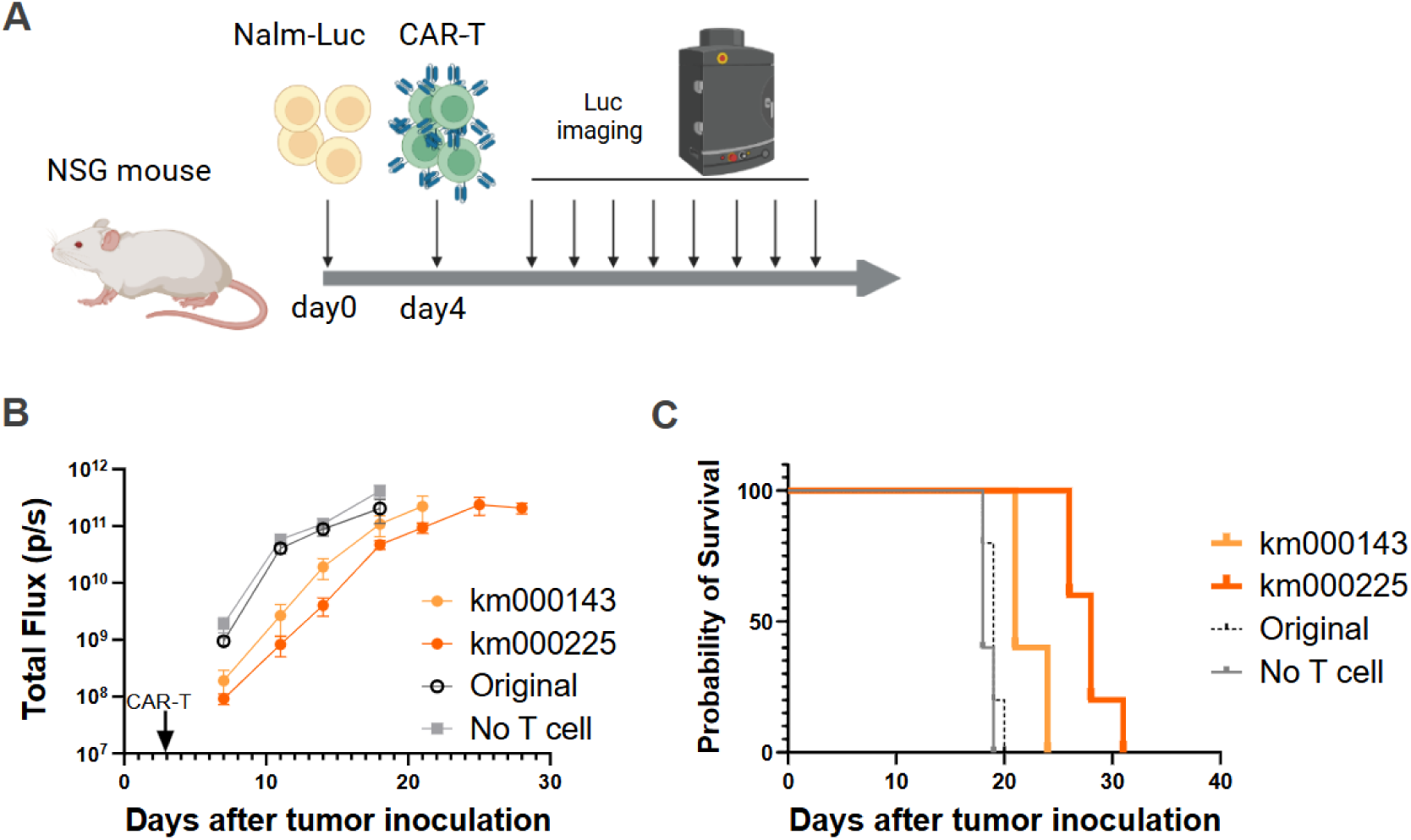
*In vivo* evaluation using PBMC-derived CAR T cells. (A) Schematic design of leukemia model experiment. Created with BioRender.com. (B) Tumor burden (total flux) quantified by photons/sec in mice treated with PBS or PBMC-derived CAR T cells at day4 (arrow). (C) Kaplan-Meyer plot indicating overall survival of mice after tumor injection

**Figure S6:**
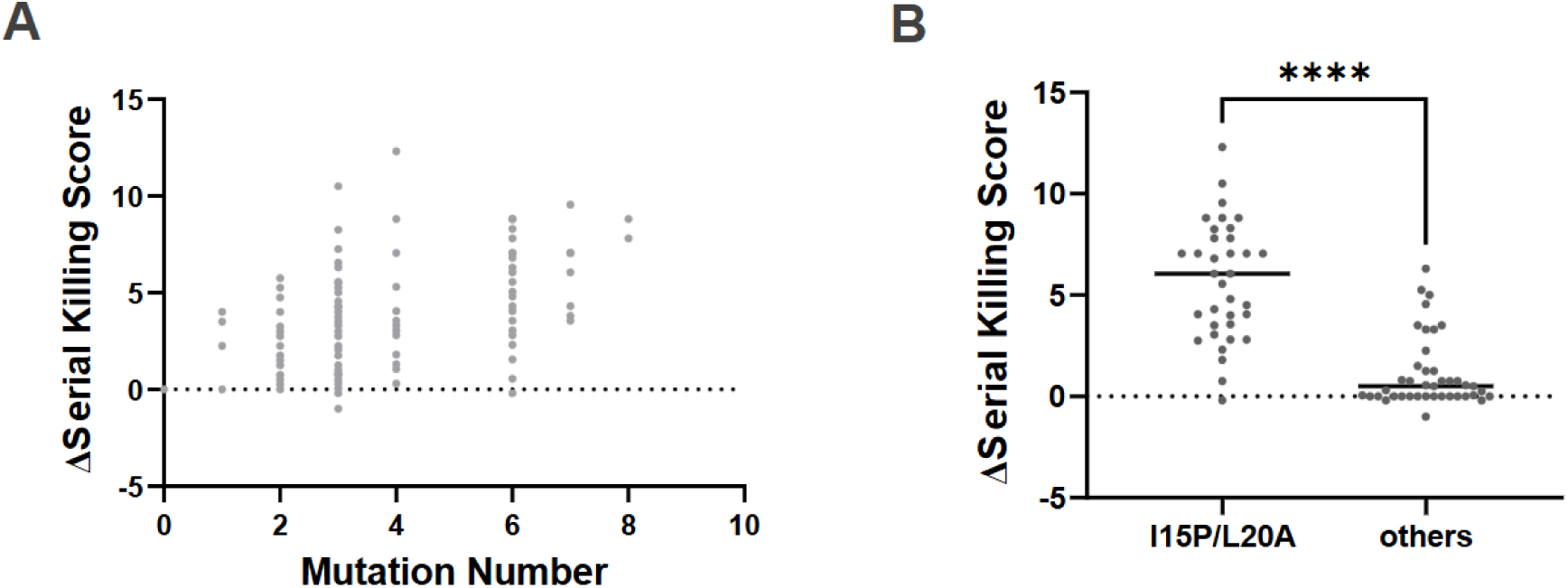
km000225-derived variants and essential combination of mutations for enhanced activity. (A) Dot plot shows the relationship between the number of mutations and Δserial killing score of km000225-derived variants. We did not test the group of variants carrying 5 mutations due to the limitation of experimental capacity. (B) Dot plot shows Δserial killing score comparing the group of variants include both I15P and L20A mutations (I15L/L20A) to the group of mutations without I15P and L20A (others). P values were determined using unpaired t test with Welch’s correction. ****, P<0.0001

